# Split RNA switch: Programmable and precise control of gene expression by ensemble of pre- and post-translational regulation

**DOI:** 10.1101/2024.09.25.614879

**Authors:** Itsuki Abe, Hirohisa Ohno, Megumi Mochizuki, Karin Hayashi, Hirohide Saito

## Abstract

Regulating gene expression in response to biomolecules is a powerful strategy for monitoring intracellular environments and controlling cellular programs. RNA switch is a synthetic mRNA-based technology that controls gene expression at the translational level in response to cellular RNA and protein molecules, thus enabling cell type-specific gene regulation and showing promise for gene therapy, regenerative medicine, and cell therapy. However, single RNA switches often lack the specificity required for practical applications due to low ON/OFF ratios and difficulty in finding distinct and single biomolecule targets. To address these issues, we developed “split RNA switches” that integrate outputs from multiple RNA switches by exploiting protein splicing, a post-translational modification mechanism. We demonstrated that split RNA switches significantly improve the ON/OFF ratio of microRNA (miRNA)-responsive ON switch systems by canceling undesirable leaky OFF level. We achieved efficient and robust target cell purification based on endogenous miRNA profiles, which was impossible with an ON switch alone. Additionally, we constructed multi-output and multi-input RNA-based synthetic circuits using split RNA switches to enable the detection of multiple miRNAs for precise gene control with logical operations. Split RNA switches illustrate a novel application of protein splicing and demonstrate the potential of post-translational processing as a comprehensive solution for advancing translational control technologies toward widespread mRNA therapies.

## Introduction

Regulating gene expression in response to biomolecules is a powerful strategy for monitoring intracellular environments and controlling cellular programs^1,2^. There are various examples of such gene-regulatory components in nature, represented by Tet repressors and riboswitches^2–4^. Researchers have repurposed such transcriptional or translational regulators to implement artificial gene regulatory systems in recent years to sense molecules of interest by introducing exogenous DNA or RNA elements into cells. Among such gene regulatory systems, translational regulation using exogenous messenger RNAs (mRNAs) is preferable for some medical applications to transcriptional regulation, which requires the introduction of foreign DNA, thus carrying a potential risk of non-specific genome integration. RNA switch is a synthetic mRNA-based technology that translationally controls gene expression in response to intracellular RNAs, proteins, and small molecules^5–11^. RNA switches, particularly those targeting endogenous microRNAs (miRNAs) or proteins distinct to certain cell types, enable the control of translation for output genes specific to these cell types. Regulating output genes such as fluorescent reporter, apoptotic, and cytotoxic genes by RNA switches enables cell type-specific classification^5,6^, cell-fate control^5,6,12,13^, and genome editing^14^. Therefore, RNA switch technology holds great potential for gene therapy, cell therapy, and regenerative medicine.

However, existing strategies using single RNA switches targeting one molecule suffer from low cell type specificity, thus hindering broad and practical applications. Target cell specificity is plagued commonly by two challenges. The first challenge is the undesired leaky expression of output protein in the OFF state (“leaky translational control”), resulting in a low ON/OFF ratio (translation level in the ON state relative to the OFF state, i.e., signal-to-noise). Even in an “ideal” situation where only the target cells in the population possess the target miRNA activity, and the non-target cells have none, leaky translation in the OFF state of RNA switch itself prevents efficient fluorescent protein-based classification or cell fate control gene-based purification of target cells based on a target biomolecule. The second challenge is the difficulty in finding a target biomolecule whose expression level in the target cell type is so different from all the other cell types that an RNA switch can clearly distinguish the cell^15^ (“lack of distinctive marker”). This issue is particularly problematic when identifying or selecting target cells from a heterogeneous population of multiple cell types. To overcome the “lack of distinctive marker” challenge, multi-input systems capable of sensing multiple molecules have been developed^7,15,16^. However, increased system complexity could generate noise, thus leading to a counterproductive situation where the overall ON/OFF ratio of the system decreases.

To address the “leaky translational control” and “lack of distinctive marker” challenges, we developed a novel mRNA-based methodology called “split RNA switch,” which employs protein splicing, a post-translational modification system, to integrate translational control by multiple RNA switches. The split RNA switch simultaneously solves these two problems by (1) improving the ON/OFF ratio by suppressing leaky output protein expression in the OFF state and (2) enabling the construction of multi-input systems. Indeed, using this system, we improved cell type specificity dramatically by increasing the ON/OFF ratio of a miRNA-responsive ON switch system from a few-fold to more than 25-fold and then achieved the first demonstration of antibiotics purification of miRNA+ target cells solely by mRNAs. Furthermore, we expanded the mechanism of leak cancellation to a dual-output system to enhance the binariness of a miRNA-responsive toggle-like system, which resulted in a clearer separation of cell types in flow cytometry two-dimensional plots. Finally, by utilizing multiple split RNA switches, each targeting different miRNAs, we constructed a two-input system capable of simultaneously sensing two types of miRNAs with logic gates.

In principle, the split RNA switch system can be adapted to any gene and implemented simply by ORF sequence design, making it directly applicable to research and medical applications using existing mRNA-based technologies (e.g., UTR design, transfection, and delivery methods), as well as circular mRNA and self-replicating RNA. The split RNA switch presents a novel strategy for orchestrating pre- and post-translational regulation, which simplifies the design of RNA-based synthetic intracellular circuits and offers a comprehensive solution for precise gene regulation using RNA switch technology.

## Results

### miRNA-responsive split ON switch system with enhanced ON/OFF ratio

MicroRNAs (miRNAs) are small noncoding RNAs that post-transcriptionally regulate gene expression of target mRNAs. With over 2600 miRNAs encoded in the human genome^17,18^ and unique expression patterns among different cell types^19–22^, miRNAs are useful markers for distinguishing cell types^15,23–27^. miRNA-sensing gene regulatory systems have been developed previously^28–30^, including miRNA-responsive RNA switches^5,6^ (Fig. 1a), thus allowing cell type specific gene regulation^5,6,12–14^. We previously developed two types of miRNA-responsive RNA switches: OFF switch^5^ and ON switch^6^. An OFF switch contains an antisense sequence (“miRNA target site”) of the target miRNA in the 5’ UTR^5^. While it is translated in the same manner as normal mRNA in cells without the target miRNA activity (miRNA-cells), translation is inhibited by miRNA binding to the target site in cells possessing the specific miRNA activity (miRNA+ cells) (Fig. 1a, left). Conversely, an ON switch has a miRNA target site downstream of the poly(A) tail, followed by an extra sequence in the 3’ terminus^6^. In miRNA-cells, translation is inhibited by the extra sequence and a miRNA target site after the poly(A) tail. In miRNA+ cells, miRNA binds to the target site, resulting in cleavage of the extra sequence^31^ and translation from the ON switch, thus functioning as a miRNA-sensing gene activator (Fig. 1a, right). However, ON switches with extra sequences often exhibit low ON/OFF ratios due to leaky OFF state translation, as shown in our prior study^6^.. Therefore, substantial efforts are still required in optimizing switch design to improve ON/OFF ratio for precise control of gene expression in a target cell.

**Fig 1.**
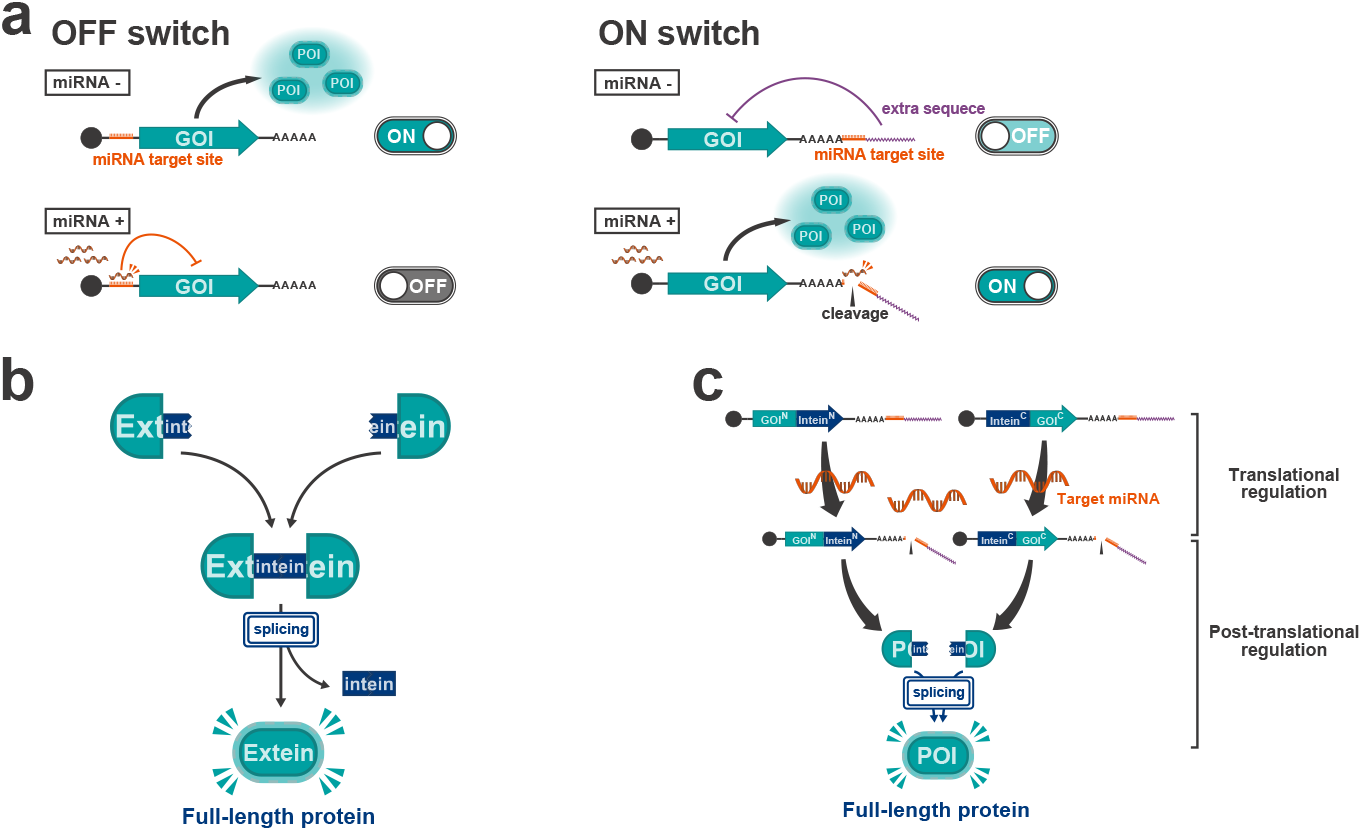
miRNA-responsive RNA switches and *trans*-protein splicing. **a**, Schematic illustration of miRNA-responsive RNA OFF and ON switches coding the gene of interest (GOI). The OFF switch (left) inhibits translation of the protein of interest (POI) in the presence of miRNA activity, while the ON switch (right) promotes translation under the same condition. **b**, Schematic illustration of trans-protein splicing. In this reaction, split-inteins linked to their flanking peptides (exteins) ligate post-translationally, excising themselves and seamlessly joining the exteins to generate a mature protein. **c**, Schematic illustration of “split ON switch,” a pair of ON switches coding protein fragments conjugated with split-inteins.

To achieve output protein control with a better ON/OFF ratio in a target cell-dependent manner without engineering the RNA switch itself, we employed protein splicing and integrated translational controls of multiple RNA switches to suppress protein activity level in the OFF state (“split RNA switch,” Fig. 2a).

**Fig 2.**
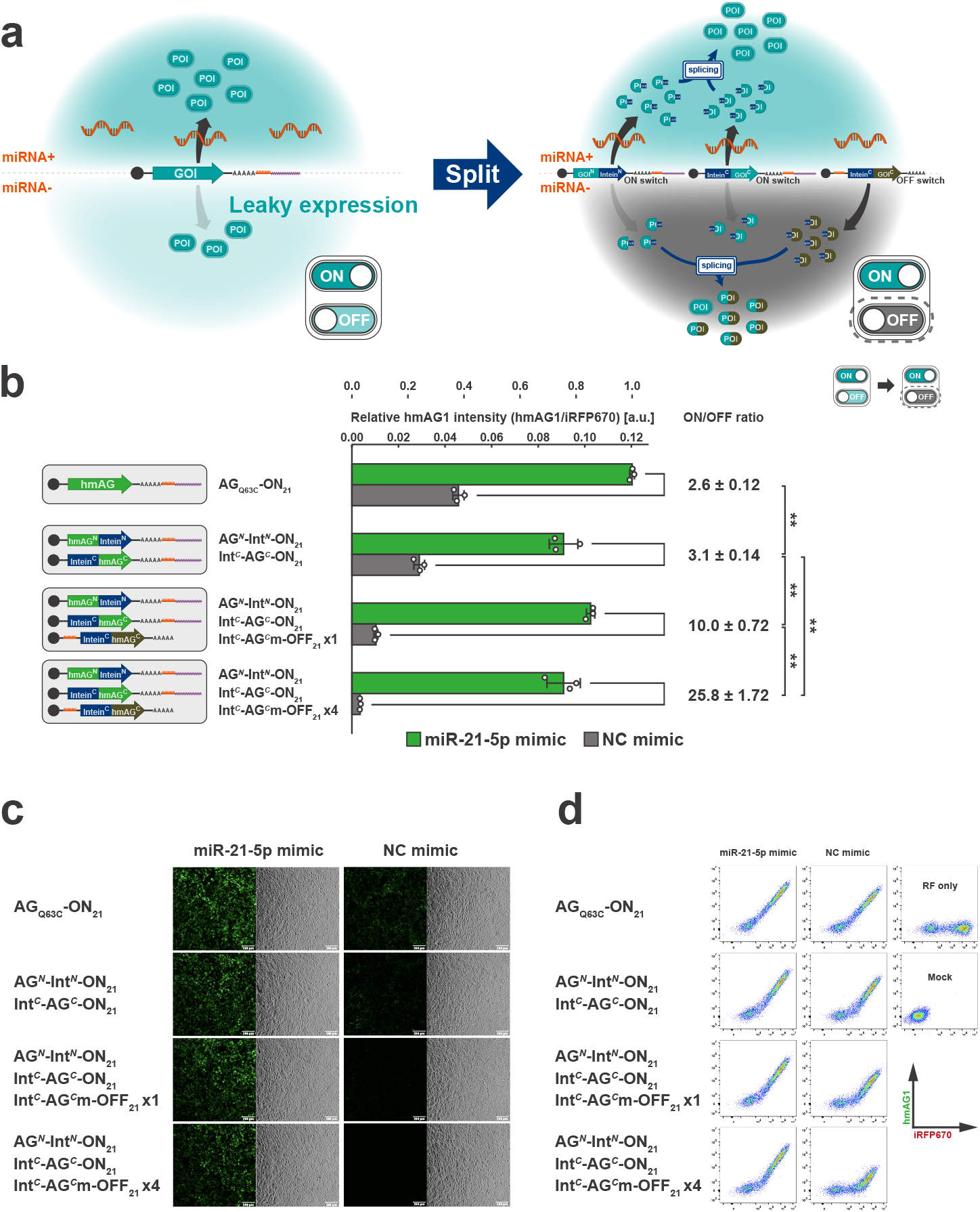
Reporter assay of split ON switch system. **a**, Schematic illustration of the strategy to improve the ON/OFF ratio of ON switch systems. Introducing a “leak-canceller”, an OFF switch coding an inactive C-terminal fragment, together with the split ON switch enables the suppression of leaky protein activity in miRNA-cells and enhances the ON/OFF ratio. **b**, Relative hmAG1 intensity (hmAG1/iRFP670) of HEK293FT cells treated with miR-21-5p or negative control (NC) mimics. Error bars represent means ± SD (n=3), and data of each biological replicate are shown as a point. Statistical analysis by two-sided Welch’s t-test, \*\**P*<0.01, n.s.: not significant (*P*>0.05). **c**, Representative microscopic images of HEK293FT cells transfected in the same conditions as shown in **b**. Green fluorescence (left) and bright-field (right) images are shown for each condition. Scale bar, 200 μm. **d**, Representative 2D flow cytometry plots. The horizontal axis shows the fluorescence intensity of iRFP670 (reference), and the vertical axis shows the fluorescence intensity of hmAG1 (reporter).

Protein-(*trans-*) splicing is a type of post-translational modification that occurs between protein fragments linked to peptide sequences called split-inteins^32,33^. Corresponding split-intein segments ligate with each other and then excise themselves from the flanking sequence called extein, seamlessly joining extein sequences, and finally generating a full-length protein^34,35^(Fig. 1b). The split-intein used in this study, Npu DnaE, requires only the N-terminal amino acid residue of the C-extein to be cysteine for efficient protein splicing^36,37^ to allow great flexibility in extein sequences, enabling reconstitution in a wide variety of proteins with a high efficiency^36,38–42^.. We designed a pair of ON switches coding protein fragments conjugated with split-inteins, termed “split ON switch” (Fig. 1c).

To improve the ON/OFF ratio of the output protein in response to target miRNA activity, we devised the split RNA switch approach based on the following mechanism (Fig. 2a). First, we introduced a pair of ON switches, coding N- and C-terminal protein fragments linked to split-inteins (split ON switch pair: GOI^N^-Intein^N^-ON and Intein^C^-GOI^C^-ON) along with an OFF switch coding an inactivated mutant C-terminal fragment linked to C-terminal split-intein (Intein^C^-GOI^C^m-OFF21, hereafter referred to as “leak-canceller”). Whereas ON switches generate N- and C-terminal fragments of the protein of interest in miRNA+ cells (Fig. 2a, upper right), little translation arose from the OFF switch. Then, the N- and C-terminal fragments are immediately spliced to form a full-length, functional protein. In contrast, in miRNA-cells (Fig. 2a, lower right), ON switches show leaky expression of N- and C-terminal fragments. Meanwhile, the OFF switch generates mutated C-terminal fragments. Because the mutated C-terminal fragments are present in larger amounts compared to the normal C-terminal fragments, most of the N-terminal fragments, leaked from the ON switch, are expected to undergo the irreversible splicing reaction with the inactivated C-terminal fragments. Therefore, the formation of full-length functional output proteins with a normal C-terminal fragment will be inhibited. As a result, the leaky activity of the output protein in miRNA-negative cells will be suppressed. Theoretically, the more OFF switches are introduced, the stronger this competitive cancellation effect will be.

To test our hypothesis, we demonstrated a reporter assay in HEK293FT cells using hmAG1 as the output protein for quantification by flow cytometry and fluorescence microscopy (Fig. 2b-d). In this reporter assay, we used a miR-21-5p mimic to induce miR-21-5p activity in HEK293FT cells (miR-21-5p activity is otherwise low in HEK293FT cells^15^), and an iRFP670-coding mRNA as a reference. In cells with a negative control (NC) mimic, the single ON switch coding full-length hmAG1(AG-ON21) exhibited over one-third of fluorescence intensity compared with cells treated with the miR-21-5p mimic, indicating substantial leaky expression in the OFF state. Similarly, a moderate ON/OFF ratio was observed when introducing only a split ON switch pair coding N- and C-terminal fragments. Meanwhile, when transfecting the split ON switch pair in equal amounts with the leak-canceller (OFF switch coding the inactivated C-terminal fragment), the fluorescence intensity in cells without miR-21-5p mimic was reduced to about 10% of the level displayed by cells with the mimic, thus increasing the ON/OFF ratio to 10-fold. Furthermore, a split ON switch pair with four times the amount of the leak-canceller resulted in an ON/OFF ratio exceeding 25 times in response to miRNA activity. The slight decrease in ON levels in the miR-21-5p mimic condition could be attributed to leaky translation from the OFF switch. From these results, we confirmed that the split ON switch system suppresses undesired leaky output in the OFF state, thereby improving the ON/OFF ratio of the ON switch system.

### Cell-fate control using drug-resistance genes and a suicide gene

Next, to verify the versatility of our split ON switch system, we applied the system to three commonly used antibiotic resistance genes-encoding mRNAs (puromycin N-acetyltransferase (PAC) (Fig. 3a), hygromycin-B 4-O-kinase (HPH) (Fig. 3c), and blasticidin-S deaminase (BSR) (Fig. 3e)) and demonstrated cell fate control based on endogenous miR-21-5p activity. In each experiment, we used a pair of split ON switches, coding N- and C-terminal fragments of the gene, and a leak-canceller, coding an inactivated C-terminal fragment of PAC. It is not necessary to prepare mutated versions of C-terminal fragments for each gene because mutated C-terminal fragments for leakage-cancellation are required only to form an inactive protein through splicing with the N-terminal fragment. We evaluated the survival rate of HeLa cells, which have high endogenous activity of miR-21-5p when cultured in the presence of antibiotics and treated with either miR-21-5p or negative control (NC) inhibitors (Fig. 3b, d, f). In all three experiments, transfection of an ON switch coding the full-length gene led to leaky survival of HeLa cells treated with miR-21-5p inhibitor and antibiotics due to translational leakage of the resistance gene. Especially in the cases of HPH and BSR ON switches, survival was nearly unchanged regardless of miR-21-5p activity. These small changes were observed presumably because the amount of transfected mRNA was so large that the quantity of leaked resistance protein was sufficient to rescue the cells almost completely. Notably, in all three experiments, the introduction of the split ON switch pair along with the leak-canceller allowed HeLa cells with high miR-21-5p activity to survive, whereas significantly suppressing the survival of HeLa cells deprived of miR-21-5p activity under antibiotics-added culture conditions. Thus, the split RNA switch system is highly versatile in terms of its output genes, which enables cell type-specific fate control using antibiotic resistance genes optimized for target cells.

**Fig 3.**
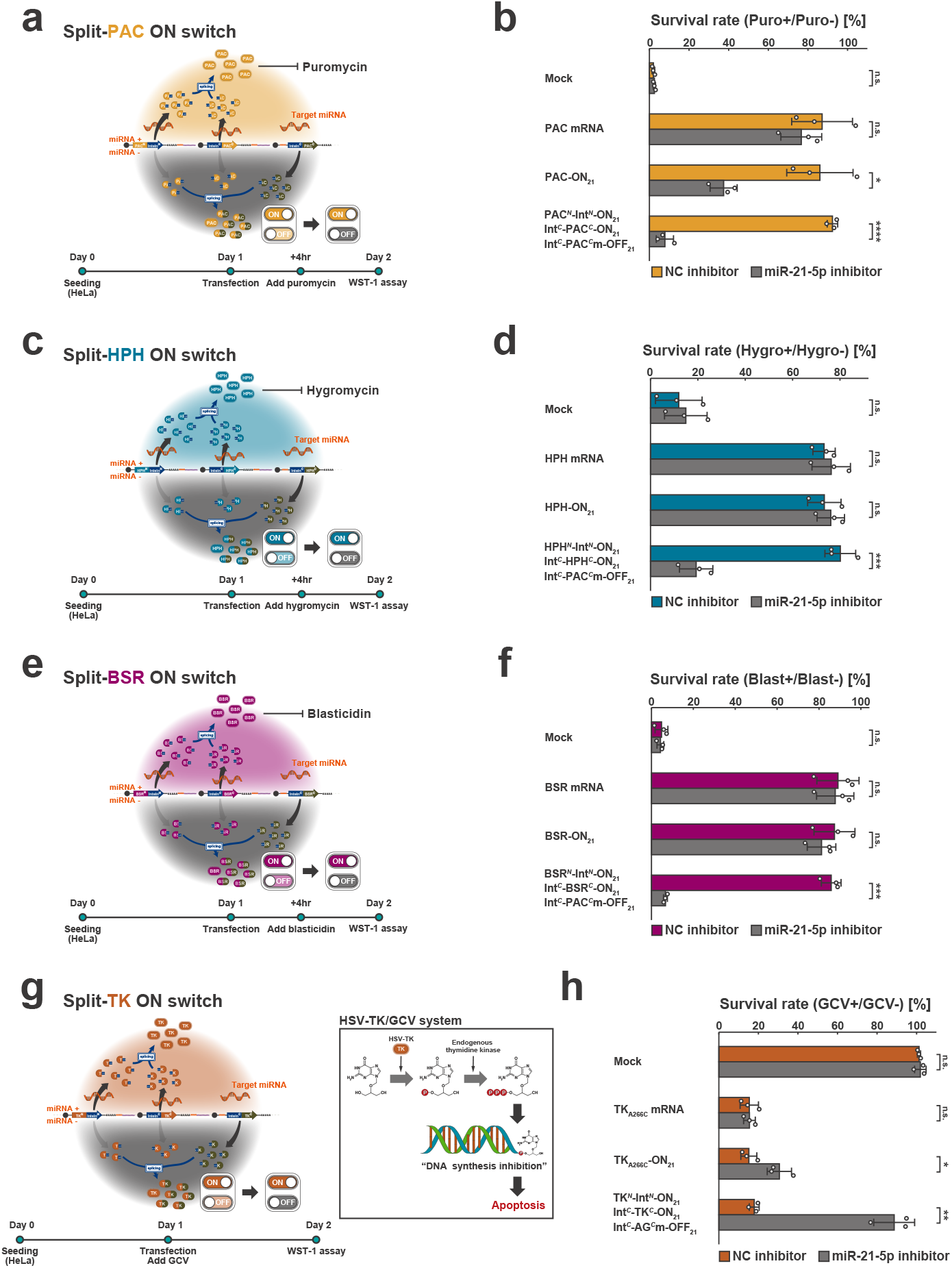
Regulation of various genes by split ON switch system. **a, c, e**, Schematic illustration of split ON switch system regulating three antibiotic resistance genes: puromycin N-acetyltransferase (PAC) (**a**), hygromycin-B 4-O-kinase (HPH) (**b**), and blasticidin-S deaminase (BSR) (**c**). The experimental time course is shown below each schematic illustration. **b, d, f**, The survival rate of HeLa cells treated with miR-21-5p or NC inhibitors as measured by the WST-1 assay. For each condition, the survival rates were calculated by dividing the viability in the presence of antibiotics by that without antibiotics. Error bars represent means ± SD (n=3). **g**, Schematic illustration of split ON switch system regulating herpes simplex virus type 1 thymidine kinase (HSV-TK). **h**, The survival rate of HeLa cells treated with miR-21-5p or NC inhibitors as measured by the WST-1 assay. For each condition, the survival rates were calculated by dividing the viability in the presence of ganciclovir (GCV) by that without GCV. Error bars represent means ± SD (n=3), and data of each biological replicate are shown as a point. Statistical analysis by two-sided Welch’s t-test (**b, d, f, h**), \**P*<0.05, #x002A;\**P*<0.01, \*\*\**P*<0.001, \*\*\*\**P*<0.0001, n.s.: not significant (*P*>0.05).

In addition to these genes for cell type-specific drug selection, we regulated herpes simplex virus 1 thymidine kinase (HSV-TK) by the split ON switch system and demonstrated target cell (miRNA+)-specific apoptosis induction that is dependent on both target miRNAs and ganciclovir (GCV)^43^ (Fig. 3g). In the HSV-TK/GCV system, ganciclovir, a thymidine analog, is monophosphorylated exclusively by the exogenous HSV-TK^44^. Then, the monophosphate is further phosphorylated by endogenous TK to produce GCV-triphosphate. This triphosphate is incorporated into replicating DNA, terminating the elongation of DNA strands and finally inducing apoptosis^45–47^. We treated HeLa cells with GCV at the same time as the transfection of RNA switches and miRNA inhibitors (miR-21-5p or NC inhibitors). After 24 hours, we evaluated cell viability by the WST-1 assay (Fig. 3h). TKA266C, a full-length HSV-TK with A266C, an expected single amino acid mutation introduced after protein splicing, reduced the survival rate of HeLa cells to about 15% in the presence of GCV compared to that without GCV. When using the ON switch coding TKA266C, the survival rate of HeLa cells with miR-21-5p inhibitor declined to 30.6 ± 6.1% by GCV treatment, presumably caused by leaky translation from the ON switch. Meanwhile, the split ON switch pair with a leak-canceller suppressed the leakage of HSV-TK activity, and cell survival remained high in HeLa cells treated with miR-21-5p inhibitor and GCV (88.7 ± 10.3%). These results indicated the utility of a split switch system across output proteins for improving the ON/OFF ratio.

### Cell type-specific selection based on endogenous miRNA activity

Next, we performed a cell type-specific selection experiment based on the differences of endogenous miRNA activity between mixed cell types using split ON switches coding BSR (Fig. 4a). In this experiment, we used HeLa cells (high miR-21-5p activity) as target cells to be selected and HEK293FT cells (low miR-21-5p activity) as non-target cells to be eliminated. To distinguish each cell type from the mixture, we used HeLa cells expressing hmAG1-M9 (HeLa-hmAG1-M9) and HEK293FT cells expressing iRFP670-M9 (HEK293FT-iRFP670-M9). After the introduction of the miR-21-5p responsive BSR-ON switch system into a mixture of HeLa and HEK293FT cells, BSR should be active specifically in HeLa cells, resulting only in HeLa cell survival in the presence of blasticidin. After seeding the mixture of HeLa-hmAG1-M9 and HEK293FT-iRFP670-M9, we transfected RNA switches and added blasticidin 24 hours later, followed by imaging and flow cytometry analysis after three days (Fig. 4b, c, d. When a single ON switch coding BSR was used, the ratio of HeLa to HEK293FT cells (58.6 ± 4.0%) was comparable to the results of untreated cells (neither transfection nor blasticidin treatment) (58.9 ± 6.6%) and cells with BSR mRNA (60.8 ± 2.2%). Meanwhile, the introduction of the split ON switch pair along with the leak-canceller suppressed HEK293FT cell viability and the purification of HeLa cells with a purity of 96.5 ± 0.61%. From these results, we concluded that the enhanced ON/OFF ratio of the split ON switch system enabled cell type-specific purification based on drug resistance, previously unachievable with a single ON switch.

**Fig 4.**
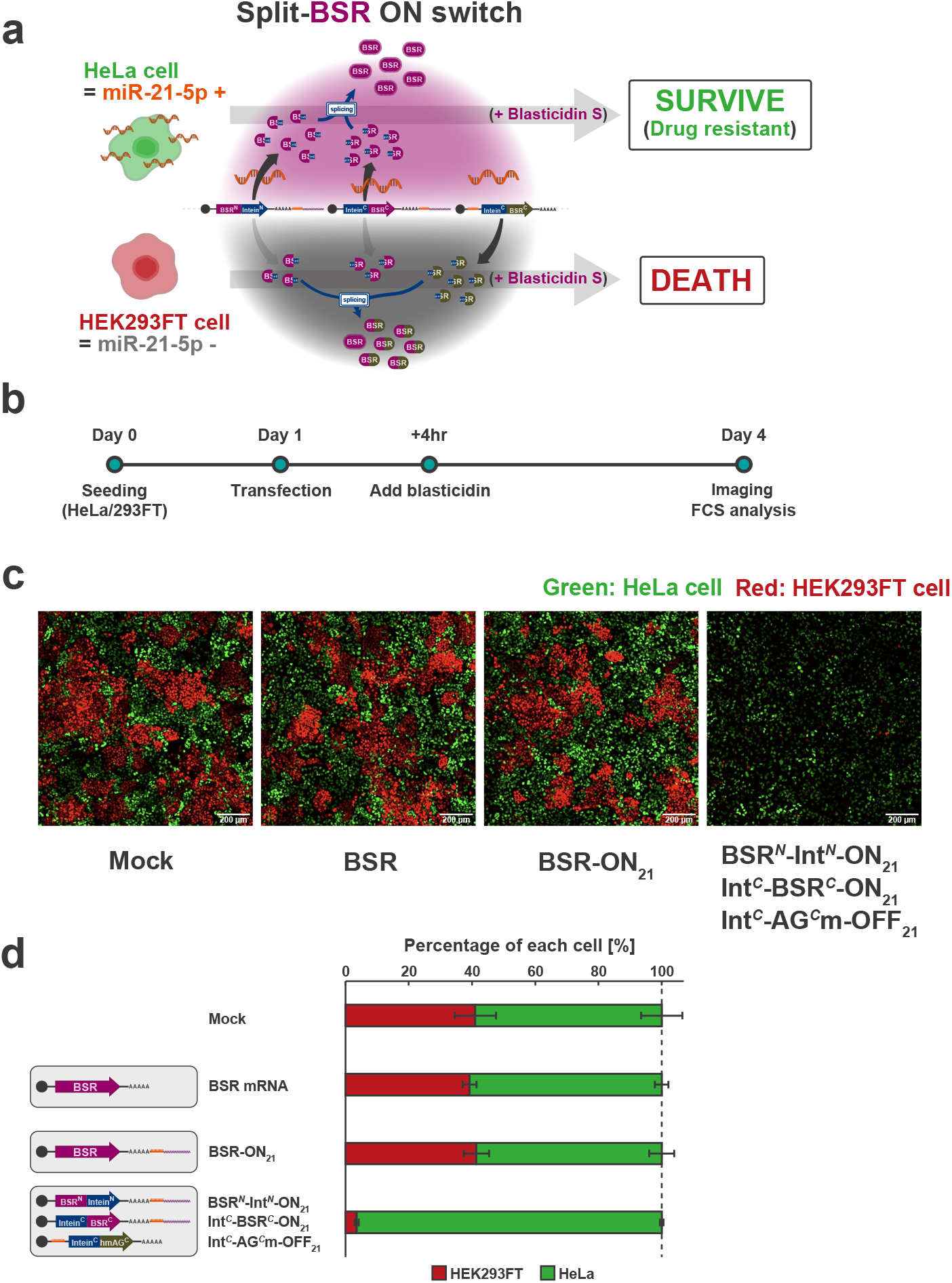
Cell type-specific selection based on endogenous miRNA activity. **a**, Schematic illustration of the miR-21-5p-responsive split ON switch system regulating blasticidin-S deaminase (BSR). This system suppresses the leaky activity of BSR in miR-21-5p negative cells (HEK293FT cells) compared to the conventional single ON switch while maintaining the drug-resistance gene in miR-21-5p positive cells (HeLa cells). **b**, Schematic illustration of the experimental procedure. **c**, Representative merged fluorescence images of the mixture of HeLa and HEK293FT cells transfected with mRNAs. HeLa cells stably expressing hmAG1-M9 (HeLa-hmAG1-M9) and HEK293FT cells stably expressing iRFP670-M9 (HEK293FT-iRFP670-M9) were used to distinguish each cell line. Scale bar, 200 μm.==**d**, Percentage of HeLa and HEK293FT cells. The number of each cell type was counted, and the percentage of each cell line was calculated. Error bars represent means ± SD (n=3).

### Improving the performance of 2-outputs with a split toggle-like system

In flow cytometry (FC) analysis, miRNA-responsive RNA switches can be a powerful tool to distinguish different cell types based on their endogenous miRNA profiles^5,12,15^. However, a single RNA switch often fails to completely separate two cell populations, mainly due to a low ON/OFF ratio of RNA switch and minute differences in endogenous miRNA activity^6,15^. In contrast, a “toggle-like system,” a two-output system switching its outputs in response to the target miRNA activity, offers clearer separation in FC 2D plots. A simple way to construct a toggle-like system using RNA switches is to introduce a pair of ON and OFF switches, each coding different fluorescent proteins, targeting the same miRNA (“normal toggle-like system”) (Fig. 5a, left). Nevertheless, in the case of the normal toggle-like system, the leaky translation from each switch could still undermine the binariness of the system.

**Fig 5.**
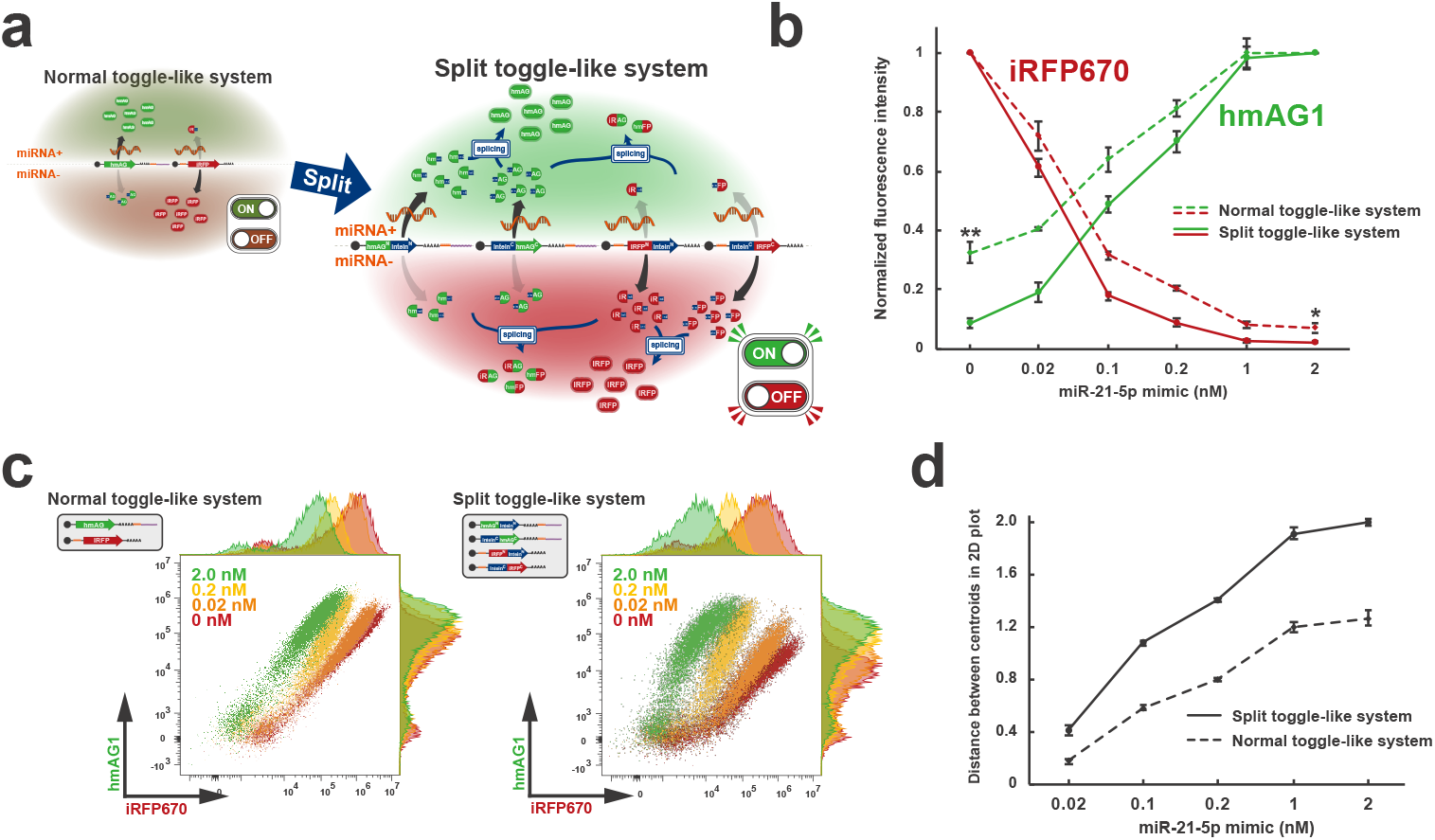
Engineering of a toggle-like system using split-intein. **a**, Schematic illustration of a toggle-like (2-output) system that displays hmAG1 fluorescence in the presence of target miRNA activity and iRFP670 fluorescence in its absence. “Normal toggle-like system” refers to a system composed of an ON switch coding full-length hmAG1 and an OFF switch coding full-length iRFP670, and “split toggle-like system” refers to a system composed of a pair of ON switches coding split hmAG1 fragments and a pair of OFF switches coding split iRFP670 fragments. **b**, Normalized hmAG1 and iRFP670 fluorescence intensity of HEK293FT cells treated with various miR-21-5p mimic concentrations. Normalized hmAG1 intensity was calculated by normalizing the intensity at 2 nM mimic concentration, and normalized iRFP670 intensity was calculated by normalizing the intensity at 0 nM mimic concentration. Error bars represent means ± SD (n=3). Statistical analysis by two-sided Welch’s t-test, \**P*<0.05, \*\**P*<0.01. **c**, Representative 2D flow cytometry plots of HEK293FT cells treated with various miR-21-5p mimic concentrations. The horizontal axis shows the fluorescence intensity of iRFP670 (reference), and the vertical axis shows the fluorescence intensity of hmAG1.==**d**, Euclidean distance between the centroid of the 0 nM mimic concentration plot and those of various mimic concentrations in the logarithmic 2D plot in **c**. Error bars represent means ± SD (n=3).

We hypothesized that the combination of protein splicing and RNA switch could enhance the binariness of a toggle-like system. As a proof-of-concept, we devised the “split toggle-like system” using two pairs of switches: two ON switches coding the N-terminal and C-terminal of hmAG1 and two OFF switches coding the N-terminal and C-terminal of iRFP670 (Fig. 5a, right). In miRNA+ cells, the high abundance of hmAG1 fragments translated from ON switches inhibits splicing between iRFP670 fragments—leaked from the OFF switches—thereby inhibiting the formation of functional iRFP670. In miRNA-cells, the opposite is expected: abundant iRFP670 fragments translated from OFF switches inhibit splicing between hmAG1 fragments leaked from the ON switches, thus inhibiting the formation of functional hmAG1. In the entire system, the binary trend, in which miRNA+ cells exhibit green fluorescence and miRNA-cells display red fluorescence, should be intensified, resulting in better separation of the two cell populations in FC 2D plots. To test the system, we transfected the four switches (AG^*N*^-Int^*N*^-ON21, Int^*C*^-AG^*C*^-ON21, RF^*N*^-Int^*N*^-OFF21, Int^*C*^-RF^*C*^-OFF21) into HEK293FT cells with varying concentrations of miR-21-5p mimic, followed by flow cytometry analysis 24 hours later (Fig. 5b, c). In the split toggle-like system, hmAG1 fluorescence intensity under the condition without miR-21-5p mimic was suppressed to 8.4 ± 1.6% of the condition with 2 nM mimic, significantly lower than that in the normal toggle-like system (32.4 ± 3.4%). Notably, iRFP670 fluorescence intensity at sufficient mimic concentrations (2 nM) was also suppressed to 2.1 ± 0.35% of the condition without mimic and significantly lower than that in the normal toggle-like system (6.9 ± 1.5%). Additionally, the distance between the centroids of the two cell populations at 0 nM mimic and each population in logarithmic 2D plots was approximately 1.6-2.3 times greater compared to the normal toggle-like system (Fig. 5d). From these results, we concluded that splitting the output genes coded in RNA switches in a toggle-like system improves the distinction of the two states, which allows for clearer classification of cell types based on differential miRNA activity.

### Construction of mRNA-based intracellular two-input logic gates

In addition to improving the ON/OFF ratio of the ON switch system and the binariness of the toggle-like system, the combination of RNA switch and protein splicing enables the construction of multi-input intracellular systems that sense multiple miRNAs to regulate output protein function. For example, introducing a pair of ON switches with different miRNA target sites, each of which codes the N- and C-terminal fragments of a gene, allows for the construction of an “AND gate,” in which the protein of interest functions only in cells that exhibit both miRNA activities (Fig. 6a).

**Fig 6.**
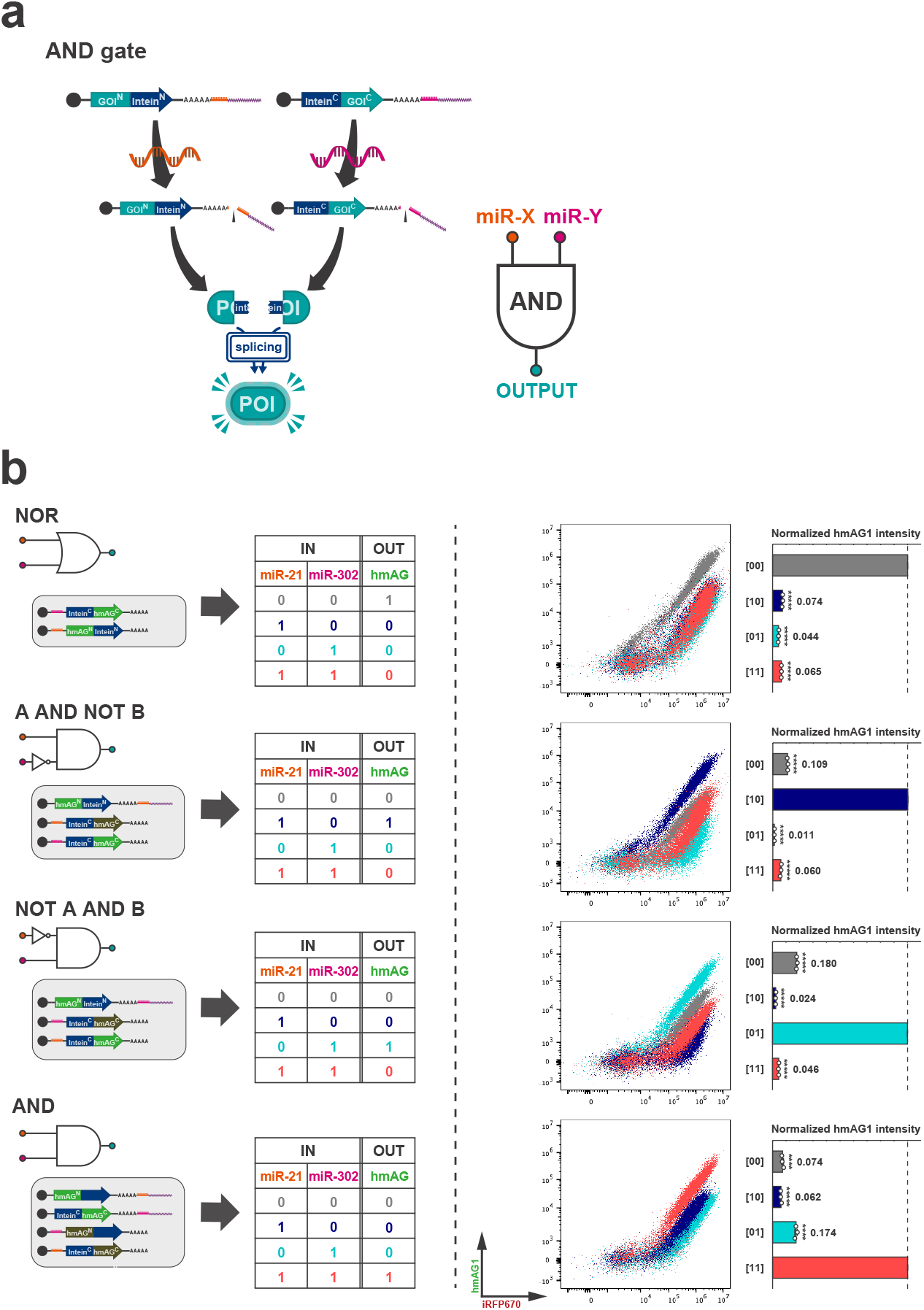
Intracellular two-input logic circuit using split-intein. **a**, Schematic illustration of a two-input logic gate using split-intein, exemplifying the AND gate, in which an output protein is activated only in the presence of both of two different target miRNAs. **b**, Four types of two-input logic circuits. Different colors correspond to those in the truth table, representative 2D flow cytometry plots, and bar graphs, respectively. MiR-21-5p and miR-302a-5p mimics were used as inputs. For example, the input pattern [10] colored in blue means miR-21-5p was present while miR-302a-5p was absent. The representative 2D flow cytometry plot for each circuit is shown as an overlay of scatter plots for all four input patterns. The relative fluorescence intensity of hmAG1 (hmAG1/iRFP670) was normalized by the highest value for each circuit ([00] in NOR, [10] in A AND NOT B, [01] in NOT A AND B, and [11] in AND). Error bars represent means ± SD (n=3), and data of each biological replicate are shown as a point. Statistical analysis by two-sided Dunnett’s test for the ON state of each circuit, \*\*\*\**P*<0.0001, \*\*\*\*\**P*<0.00001.

As a proof-of-concept experiment, we constructed four types of two-input logic gates (NOR, A AND NOT B, NOT A AND B, and AND) in HEK293FT cells treated with different combinations of miR-21-5p or miR-302a-5p mimics. After introducing ON or OFF switches responsive to miR-21-5p or miR-302a-5p, we evaluated their performance in a reporter assay using hmAG1 as the output (Fig. 6b). Each system is expected to exhibit hmAG1 fluorescence activity only in cells with specific combinations of miR-21-5p and miR-302a-5p activities (denoted as [00], [10], [01], [11]). In the three systems (A AND NOT B, NOT A AND B, and AND) in which we used ON switches, we added a leak-canceller, an OFF switch coding an inactive, opposing fragment with the same miRNA target site as the corresponding ON switch, to prevent translational leakage from ON switches. In the reporter assay, the values of the OFF state were suppressed to less than 10% of the ON state in the NOR gate and less than 20% of the ON state in the A AND NOT B, NOT A AND B, and AND gates. FC 2D plots indicated that all the logic gates regulated hmAG1 dramatically enough to distinguish a cell subpopulation from other states to near completion in the scatter plots (Fig. 6b, right). Considering the mechanisms of each logic gate, increasing the amount of leak-canceller can further suppress the leakage in OFF states. These results show that the integration of translational control by multiple RNA switches via protein splicing simultaneously enables the construction of RNA-based intracellular two-input systems and the suppression of leakage in their OFF states.

## Discussion

RNA switch is an RNA-based translational control tool that holds great potential for regenerative medicine and mRNA therapeutics to enable cell type-specific gene regulation without the risk of harmful genomic mutagenesis. While prior studies have shown the impact of RNA switch, small ON/OFF ratios of output and difficulties in finding optimal target candidate molecules have often prevented practical applications such as cell classification and purification. In this study, to address these issues, we have designed “split RNA switches” that utilize protein splicing, a type of post-translational modification. The combination of RNA switch and protein splicing allows the integration of translational outputs from multiple RNA switches at the post-translational level, thereby achieving high cell type specificity. We designed the split RNA switches, RNA switches coding split genes conjugated by split-intein sequences (Fig. 1). By using a set of split RNA switches coding functional or non-functional protein fragments, we implemented a miRNA-responsive ON switch system in HEK293 cells with over 25-fold ON/OFF ratio, resulting from suppression of undesired leaky output in OFF state (Fig. 2). Results from the hmAG1 reporter assay implied that increasing the amount of OFF switch could enable further suppression of the leakage in the OFF state. Although we used inactive C-terminal fragments with a single amino acid mutation as the output protein for the leak-canceller in this research, it would be unnecessary to prepare mutated C-terminal fragments for each gene because mutated C-terminal fragments for leak-canceller are only required to form an inactive protein through splicing with the N-terminal fragment. Considering this point, the extein of the leak-canceller should work with a shorter length, and thus smaller volumes of leak-canceller mRNA could enable the same level of leak-canceling efficiency as in this study.

We demonstrated efficient cell type-specific cell fate control with split ON switch systems outputting three types of antibiotic-resistant genes (Fig. 3). In these experiments, we successfully reduced OFF-target cell survival. Additionally, we confirmed that the split-BSR ON switch system with a leak-canceller allowed for the efficient purification of HeLa cells from a mixture with HEK293FT cells based on the difference of endogenous miRNA activity (Fig. 4). In the mock condition of the cell purification experiment, the final proportion of HEK293FT cells had increased to more than 40% of the total after five days, despite an initial seeding at a HeLa-AG:293FT-RF ratio of 5:1. This result indicates that HEK293FT cells are less sensitive to blasticidin-dependent apoptotic pathways and more resistant to cell death than HeLa cells. The split BSR-ON switch system with leak-canceller achieved a high ON/OFF ratio, sufficient to purify HeLa cells beyond the difference of blasticidin-sensitivity of cell types. Furthermore, we demonstrated the activation of HSV-TK in a manner dependent on the target miRNA and GCV (Fig. 3) to explore the application of the split-ON switch system in medical scenarios, inducing cell-specific apoptosis and eliminating unwanted cells without direct off-target effects in the absence of GCV. These results indicate that split RNA switches can provide precise cell fate control, which plays a pivotal role in cell therapy and regenerative medicine, by simply introducing mRNAs. Moreover, the versatility in the output genes observed in this methodology can be leveraged for various medical applications, including cell/tissue-specific gene therapy. A major limitation of the CRISPR-Cas9 system in gene therapy is the size of Cas9, which hinders efficient delivery via recombinant adeno-associated viruses (rAAV)^49,50^. To address this issue, gene regulation by split Cas proteins and optimal split-site identification in Cas proteins have been demonstrated in previous studies^39,51^. The split RNA switch system can be directly applied to genome editing technologies using split Cas9 to improve the specificity of cell type-specific genome editing.

In addition to the “single-input single-output” systems, we designed a two-output system in which each of the two types of fluorescent proteins cancels the fluorescence of the other. The split toggle-like system separated cell subpopulations with different miRNA levels more clearly than using pairs of ON/OFF switches coding full-length proteins (Fig. 5). The leak-canceller used in this experiment, coding an inactive C-terminal fragment, resulted in inactive canceling products by splicing with cognate N-terminal fragments. By contrast, engineering the canceling products to have functional properties could be an alternative strategy for implementing two-output systems. For example, consider the blue fluorescent protein (BFP) and green fluorescent protein (GFP), which differ by a single amino acid^52^. The introduction of normal mRNA for the N-terminal fragment of GFP, along with an ON switch for the GFP C-terminal fragment and an OFF switch for the BFP C-terminal fragment, could result in green fluorescence in miRNA+ cells and blue fluorescence in miRNA-cells, with the green fluorescence effectively quenched. A more sensitive two-output system could identify and isolate a specific cell type based on the miRNA profile from highly heterogeneous cell populations such as hematopoietic lineages, in which miRNAs act as master regulators of transcriptional programs, and their expression patterns are known to reflect biological relationships among them^22,53^.

Moreover, we demonstrated four types of intracellular two-input logic gates with split RNA switches (Fig. 6). We confirmed that these two-input logic gates output high protein activity in cells with a certain combination of two types of miRNA activities but not in other conditions. In addition to the Npu DnaE split intein used in this study, there are different types of inteins with splicing-orthogonality to each other. By using split gene fragments with the multiple orthogonal inteins, we could design multi-input systems that regulate the output gene in response to more than three types of miRNAs^54^. RNA-binding proteins (RBPs) have often been incorporated into RNA-based synthetic circuits in previous studies^7,16,55^. However, overexpression of widely used RBPs such as L7Ae has been reported to increase the complexity of gene networks and cause cytotoxicity^56^. The multi-input systems demonstrated in this study do not require RBPs for their construction, thus making them simpler and safer to use than previous systems.

One problem with the split ON switch is that the increased amount of OFF switches introduced potentially leads to an undesired decrease in the ON level (activity of the output protein in the presence of target miRNA activity) due to leakage from OFF switches. The decreased ON level observed in Fig. 2 is likely attributable to this effect. Additionally, competitive inhibition by inactive C-terminal fragments, produced from the leak-cancellers used in this study, is likely less efficient in OFF level suppression compared to monomeric (1:1) inhibition, as seen in the Barnase-Barster system^6^. This challenge could be improved by engineering the leak-canceller itself. Engineering of the intein sequence on the leak-canceller side^57^ or the extein sequence adjacent to intein^37,58^ has been reported to enhance the splicing efficiency in previous studies. The appropriate sequence design could enable higher leak-cancellation efficiency with only small amounts of leak-cancellers introduced.

In conclusion, we report that trans-protein splicing facilitated by split-inteins allows for the integration of outputs from RNA-based translational control systems, thereby enabling the generation of more desirable cellular outcomes. To our knowledge, this is the first demonstration of combining mRNA-based translational control with post-translational processing. The range of potential applications for “split RNA switches” is not limited to miRNA-responsive elements but could be readily extended to protein-responsive and other small molecule-responsive RNA switches^59^. It would be possible to construct intracellular logic gates that regulate gene expression in response to certain combinations of different types of molecules. Even if future optimization of individual RNA switches can improve their performance, further enhancement in their specificity may arise from adopting the split RNA switch approach presented here. Split RNA switch not only represents a novel application of protein splicing but also demonstrates the potential of post-translational processing as a comprehensive solution for existing translational control technologies toward practical application.

## Method

### Preparation of modified mRNA

The template DNA for in vitro transcription was made by PCR with KOD One (TOYOBO). All mRNAs were generated using the above PCR products and MEGAscript T7 Kit (Ambion). To suppress immune responses and enhance translation efficiency, we used *N*1-methylpseudouridine-5′-triphosphate, mΨ, instead of uridine-triphosphate, U, and Cap Analog, CleanCap AG or CleanCap AG (3’ OMe) (TriLink). Reaction mixtures were incubated at 37 °C for up to 4 h, mixed with TURBO DNase (Ambion), and further incubated at 37 °C for 30 min to remove the template DNA. The resulting mRNAs were purified using Monarch RNA Cleanup Kit (New England Biolabs), incubated with Antarctic Phosphatase (New England Biolabs) at 37 °C for 30 min, and then purified again using the same kit.

### Synthetic miRNA mimics and inhibitors

MiRNA mimics are small, chemically modified double-stranded RNAs that mimic endogenous miRNAs. mirVana miRNA Mimics (hsa-miR-21-5p, hsa-miR-302a-5p, and Negative Control #1) (Thermo Fisher Scientific) were used as mimic molecules in HEK293FT cells. The negative control mimic has a random sequence validated to have no effect on endogenous mRNAs.

MiRNA inhibitors are chemically modified, single-stranded oligonucleotides that bind specifically to and inhibit endogenous target miRNAs. mirVana miRNA Inhibitors (hsa-miR-21-5p and Negative Control #1) (Thermo Fisher Scientific) were used as the miRNA inhibitors in HeLa cells.

### Culture condition

HeLa cells and HeLa cells stably expressing hmAG1 containing a nuclear localization signal M9 (HeLa-hmAG1-M9) were cultured in DMEM High Glucose medium (Nacalai Tesque) supplemented with 10% FBS (Biosera). HEK293FT cells and HEK293FT cells stably expressing iRFP670 containing M9 (HEK293FT-iRFP670-M9) were cultured in DMEM High Glucose medium supplemented with 10% FBS, 1× MEM Non-Essential Amino Acids Solution (Thermo Fisher Scientific), 1 mM sodium pyruvate (Sigma Aldrich), and 2 mM L-glutamine (Thermo Fisher Scientific). HeLa and HEK293FT cells were harvested and seeded as follows: after a rinse with PBS, cells were incubated with Trypsin-EDTA (0.25%) (Thermo Fisher Scientific) for 5 min at 37 °C. After being suspended in culture medium, the necessary number of cells were dispensed for seeding..

### mRNA transfection

All transfections were performed using Lipofectamine MessengerMAX (Thermo Fisher Scientific) according to the manufacturer’s protocol. The transfection reagent was applied at scales of 50 μl and 10 μl for experiments in 24-well and 96-well plates, respectively. Opti-MEM (Thermo Fisher Scientific) was used as a buffer for MessengerMAX. The MessengerMAX reagent and Opti-MEM were mixed for 10 min. mRNAs with or without miRNA mimics or miRNA inhibitors diluted with Opti-MEM were mixed with the above reagent for 5 min before the mixtures were added to culture medium.

### Imaging and image processing

Cell images were captured using CellVoyager CQ1 (Yokogawa Electric Corporation). Image processing was performed using an ImageJ plugin (https://github.com/yfujita-skgcat/image_converter).

### Flow cytometry analysis (24-well plate)

After a wash with 100 μl of PBS, cells were incubated with 100 μl of Trypsin-EDTA (0.25%) for 5 min at 37 °C, and 150 μl of culture medium was added. The cell suspensions were analyzed using CytoFLEX (Beckman Coulter) and 525/40 nm Bandpass with OD1 Filter for hmAG1 and 660/10 nm Bandpass Filter for iRFP670. Data was extracted from the Flow Cytometry Standard (FCS) files and analyzed using FlowJo 10.8.2 software.

In the case of 96-well plates, after a rinse with 100 μl of PBS, cells are treated with 50 μl of Trypsin-EDTA (0.25%), followed by the addition of 100 μl of culture medium to collect the cell suspension.

### Cell proliferation assay (WST-1 assay)

Medium supplemented with 1/10 of WST-1 reagent (Roche Diagnostics KK) was prepared and used to replace the medium of the transfected cells. After incubation at 37°C for 1 h, absorbance was measured at 440 nm (Measurement wavelength) and 620 nm (Reference wavelength) by Spark (Tecan). The data were subtracted by the value of blanks and normalized by the results of untreated mock-transfected samples.

### Reporter assay

On the day before transfection, HEK293FT cells were seeded onto 24-well plates. At 24 h after seeding, mRNAs and miRNA mimics were transfected by Lipofectamine MessengerMAX reagent at a 50-μl scale. Fluorescence images of the cells were captured, and the samples were analyzed by flow cytometry 24 h after the transfection.

### Evaluation of split ON switch coding HSV-TK

HeLa cells were seeded in 96-well plates 24 h before transfection of mRNAs and miRNA inhibitors. At the same time as transfection, the medium was replaced with medium containing ganciclovir. At 24 h after the transfection, the cells were analyzed by WST-1 assay.

### Evaluation of split ON switch coding antibiotics resistance genes

HeLa cells were seeded in 96-well plates 24 h before transfection of mRNAs and miRNA inhibitors. At 4 h after transfection, culture medium was changed to containing puromycin (Invivogen), blasticidin S (Invivogen), or hygromycin B (Nacalai Tesque). At 24 h after the transfection, the cells were analyzed by WST-1 assay.

### HeLa cell selection from HeLa and HEK293FT cell coculture

HeLa-hmAG1-M9 (HeLa-AG) and HEK293FT-iRFP670-M9 (293FT-RF) were seeded in 96-well plates (ratio, HeLa-AG:293FT-RF = 5:1) 24 h before mRNAs transfection. Mixtures of HeLa-AG and 293FT-RF were cultured in the medium for the HEK293FT cells. At 4 h after transfection, culture medium was changed to containing blasticidin S. On day 4, fluorescence images of the cells were captured, and the samples were analyzed by flow cytometry.

## Acknowledgements

We thank Dr. Kelvin K. Hui (Kyoto University) for proofreading the manuscript. We also thank Dr. Yoshihiko Fujita (Kyoto University) for giving us HeLa-hmAG1-M9 and HEK293FT-iRFP670-M9. This work was supported by the JSPS KAKENHI (Grant Numbers JP20H05626, JP20H05701, JP20K12644), Japan Agency for Medical Research and Development (AMED) (22bm0104001, 23bm1223002h0002), iPS Cell Research Fund from Center for iPS Cell Research and Application (Kyoto University), ISHIZUE 2023 of Kyoto University, and Inamori Research Grants from Inamori Foundation.

## Reference

1. Hossain, G. S., Saini, M., Miyake, R., Ling, H. & Chang, M. W. Genetic Biosensor Design for Natural Product Biosynthesis in Microorganisms. Trends Biotechnol. 38, 797–810 (2020).

2. Alexandrov, K. & Vickers, C. E. In vivo protein-based biosensors: seeing metabolism in real time. Trends Biotechnol. 41, 19–26 (2023).

3. Gossen, M. et al. Transcriptional activation by tetracyclines in mammalian cells. Science 268, 1766–1769 (1995).

4. Kavita, K. & Breaker, R. R. Discovering riboswitches: the past and the future. Trends Biochem. Sci. 48, 119–141 (2023).

5. Miki, K. et al. Efficient Detection and Purification of Cell Populations Using Synthetic MicroRNA Switches. Cell Stem Cell 16, 699–711 (2015).

6. Fujita, Y. et al. A versatile and robust cell purification system with an RNA-only circuit composed of microRNA-responsive ON and OFF switches. Sci Adv 8, eabj1793 (2022).

7. Wroblewska, L. et al. Mammalian synthetic circuits with RNA binding proteins for RNA-only delivery. Nat. Biotechnol. 33, 839–841 (2015).

8. Kawasaki, S., Fujita, Y., Nagaike, T., Tomita, K. & Saito, H. Synthetic mRNA devices that detect endogenous proteins and distinguish mammalian cells. Nucleic Acids Res. 45, e117 (2017).

9. Cao, Y. et al. RNA-based translation activators for targeted gene upregulation. Nat. Commun. 14, 6827 (2023).

10. Hanson, S., Berthelot, K., Fink, B., McCarthy, J. E. G. & Suess, B. Tetracycline-aptamer-mediated translational regulation in yeast. Mol. Microbiol. 49, 1627–1637 (2003).

11. Weigand, J. E. et al. Screening for engineered neomycin riboswitches that control translation initiation. RNA 14, 89–97 (2008).

12. Parr, C. J. C. et al. MicroRNA-302 switch to identify and eliminate undifferentiated human pluripotent stem cells. Sci. Rep. 6, 32532 (2016).

13. Tsujisaka, Y. et al. Purification of human iPSC-derived cells at large scale using microRNA switch and magnetic-activated cell sorting. Stem Cell Reports 17, 1772–1785 (2022).

14. Hirosawa, M. et al. Cell-type-specific genome editing with a microRNA-responsive CRISPR-Cas9 switch. Nucleic Acids Res. 45, e118 (2017).

15. Endo, K., Hayashi, K. & Saito, H. High-resolution Identification and Separation of Living Cell Types by Multiple microRNA-responsive Synthetic mRNAs. Sci. Rep. 6, 21991 (2016).

16. Matsuura, S. et al. Synthetic RNA-based logic computation in mammalian cells. Nat. Commun. 9, 4847 (2018).

17. Griffiths-Jones, S., Grocock, R. J., van Dongen, S., Bateman, A. & Enright, A. J. miRBase: microRNA sequences, targets and gene nomenclature. Nucleic Acids Res. 34, D140–4 (2006).

18. Kozomara, A., Birgaoanu, M. & Griffiths-Jones, S. miRBase: from microRNA sequences to function. Nucleic Acids Res. 47, D155–D162 (2019).

19. Bartel, D. P. MicroRNAs: genomics, biogenesis, mechanism, and function. Cell 116, 281–297 (2004).

20. Kosik, K. S. MicroRNAs and cellular phenotypy. Cell 143, 21–26 (2010).

21. Lagos-Quintana, M. et al. Identification of tissue-specific microRNAs from mouse. Curr. Biol. 12, 735–739 (2002).

22. Chen, C.-Z., Li, L., Lodish, H. F. & Bartel, D. P. MicroRNAs modulate hematopoietic lineage differentiation. Science 303, 83–86 (2004).

23. Mansfield, J. H. et al. MicroRNA-responsive ‘sensor’ transgenes uncover Hox-like and other developmentally regulated patterns of vertebrate microRNA expression. Nat. Genet. 36, 1079–1083 (2004).

24. Brown, B. D., Venneri, M. A., Zingale, A., Sergi Sergi, L. & Naldini, L. Endogenous microRNA regulation suppresses transgene expression in hematopoietic lineages and enables stable gene transfer. Nat. Med. 12, 585–591 (2006).

25. Gentner, B. et al. Identification of Hematopoietic Stem Cell–Specific miRNAs Enables Gene Therapy of Globoid Cell Leukodystrophy. Sci. Transl. Med. 2, 58ra84–58ra84 (2010).

26. Nakanishi, H. et al. Monitoring and visualizing microRNA dynamics during live cell differentiation using microRNA-responsive non-viral reporter vectors. Biomaterials 128, 121–135 (2017).

27. Endo, K., Hayashi, K. & Saito, H. Numerical operations in living cells by programmable RNA devices. Sci Adv 5, eaax0835 (2019).

28. Rinaudo, K. et al. A universal RNAi-based logic evaluator that operates in mammalian cells. Nat. Biotechnol. 25, 795–801 (2007).

29. Brown, B. D. et al. Endogenous microRNA can be broadly exploited to regulate transgene expression according to tissue, lineage and differentiation state. Nat. Biotechnol. 25, 1457–1467 (2007).

30. Xie, Z., Wroblewska, L., Prochazka, L., Weiss, R. & Benenson, Y. Multi-input RNAi-based logic circuit for identification of specific cancer cells. Science 333, 1307–1311 (2011).

31. Pratt, A. J. & MacRae, I. J. The RNA-induced Silencing Complex: A Versatile Gene-silencing Machine *. J. Biol. Chem. 284, 17897–17901 (2009).

32. Hirata, R. et al. Molecular structure of a gene, VMA1, encoding the catalytic subunit of H(+)-translocating adenosine triphosphatase from vacuolar membranes of Saccharomyces cerevisiae. J. Biol. Chem. 265, 6726–6733 (1990).

33. Kane, P. M. et al. Protein Splicing Converts the Yeast TFP1 Gene Product to the 69-kdDSubunit of the Vacuolar H+-Adenosine Triphosphatase. Science 250, 651–657 (1990).

34. Zettler, J., Schütz, V. & Mootz, H. D. The naturally split Npu DnaE intein exhibits an extraordinarily high rate in the protein trans-splicing reaction. FEBS Lett. 583, 909–914 (2009).

35. Iwai, H., Züger, S., Jin, J. & Tam, P.-H. Highly efficient protein trans-splicing by a naturally split DnaE intein from Nostoc punctiforme. FEBS Lett. 580, 1853–1858 (2006).

36. Jillette, N., Du, M., Zhu, J. J., Cardoz, P. & Cheng, A. W. Split selectable markers. Nat. Commun. 10, 1–8 (2019).

37. Cheriyan, M., Pedamallu, C. S., Tori, K. & Perler, F. Faster protein splicing with the Nostoc punctiforme DnaE intein using non-native extein residues. J. Biol. Chem. 288, 6202–6211 (2013).

38. Siu, K.-H. & Chen, W. Control of the Yeast Mating Pathway by Reconstitution of Functional α-Factor Using Split Intein-Catalyzed Reactions. ACS Synth. Biol. 6, 1453–1460 (2017).

39. Truong, D.-J. J. et al. Development of an intein-mediated split-Cas9 system for gene therapy. Nucleic Acids Res. 43, 6450–6458 (2015).

40. Yang, J., Fox, G. C. & Henry-Smith, T. V. Intein-mediated assembly of a functional β-glucuronidase in transgenic plants. Proceedings of the National Academy of Sciences 100, 3513–3518 (2003).

41. Ho, T. Y. H. et al. A systematic approach to inserting split inteins for Boolean logic gate engineering and basal activity reduction. Nat. Commun. 12, 2200 (2021).

42. Palanisamy, N. et al. Split intein-mediated selection of cells containing two plasmids using a single antibiotic. Nat. Commun. 10, 1–15 (2019).

43. Wang, J., Lu, X.-X., Chen, D.-Z., Li, S.-F. & Zhang, L.-S. Herpes simplex virus thymidine kinase and ganciclovir suicide gene therapy for human pancreatic cancer. World J. Gastroenterol. 10, 400–403 (2004).

44. Elion, G. B. et al. Selectivity of action of an antiherpetic agent, 9-(2-hydroxyethoxymethyl) guanine. Proc. Natl. Acad. Sci. U. S. A. 74, 5716–5720 (1977).

45. Matthews, T. & Boehme, R. Antiviral Activity and Mechanism of Action of Ganciclovir. Rev. Infect. Dis. 10, S490–S494 (1988).

46. Wei, S.-J. et al. S- and G2-phase Cell Cycle Arrests and Apoptosis Induced by Ganciclovir in Murine Melanoma Cells Transduced with Herpes Simplex Virus Thymidine Kinase. Exp. Cell Res. 241, 66–75 (1998).

47. Craperi, D. et al. Increased bax expression is associated with cell death induced by ganciclovir in a herpes thymidine kinase gene-expressing glioma cell line. Hum. Gene Ther. 10, 679–688 (1999).

48. Tian, W. et al. High-throughput functional microRNAs profiling by recombinant AAV-based microRNA sensor arrays. PLoS One 7, e29551 (2012).

49. Zinn, E. & Vandenberghe, L. H. Adeno-associated virus: fit to serve. Curr. Opin. Virol. 8, 90–97 (2014).

50. Li, J., Sun, W., Wang, B., Xiao, X. & Liu, X.-Q. Protein trans-splicing as a means for viral vector-mediated in vivo gene therapy. Hum. Gene Ther. 19, 958–964 (2008).

51. Yuan, G. et al. An Intein-Mediated Split–nCas9 System for Base Editing in Plants. ACS Synth. Biol. 11, 2513–2517 (2022).

52. Heim, R., Prasher, D. C. & Tsien, R. Y. Wavelength mutations and posttranslational autoxidation of green fluorescent protein. Proc. Natl. Acad. Sci. U. S. A. 91, 12501–12504 (1994).

53. Petriv, O. I. et al. Comprehensive microRNA expression profiling of the hematopoietic hierarchy. Proc. Natl. Acad. Sci. U. S. A. 107, 15443–15448 (2010).

54. Pinto, F., Thornton, E. L. & Wang, B. An expanded library of orthogonal split inteins enables modular multipeptide assemblies. Nat. Commun. 11, 1–15 (2020).

55. Ausländer, S., Ausländer, D., Müller, M., Wieland, M. & Fussenegger, M. Programmable single-cell mammalian biocomputers. Nature 487, 123–127 (2012).

56. Wagner, T. E. et al. Small-molecule-based regulation of RNA-delivered circuits in mammalian cells. Nat. Chem. Biol. 14, 1043–1050 (2018).

57. Stevens, A. J. et al. Design of a Split Intein with Exceptional Protein Splicing Activity. J. Am. Chem. Soc. 138, 2162–2165 (2016).

58. Shah, N. H., Eryilmaz, E., Cowburn, D. & Muir, T. W. Extein Residues Play an Intimate Role in the Rate-Limiting Step of Protein Trans-Splicing. J. Am. Chem. Soc. 135, 5839–5847 (2013).

59. Ono, H. & Saito, H. Sensing intracellular signatures with synthetic mRNAs. RNA Biol. 20, 588–602 (2023).

